# Neurofeedback fMRI in the motor system elicits bi-directional changes in activity and white-matter structure in the healthy adult human brain

**DOI:** 10.1101/2020.08.06.234526

**Authors:** Cassandra Sampaio-Baptista, Heather F. Neyedli, Zeena-Britt Sanders, Kata Diosi, David Havard, YunYing Huang, Jesper L. R. Andersson, Michael Lührs, Rainer Goebel, Heidi Johansen-Berg

## Abstract

Neurofeedback can be used to alter brain activity and is therefore an attractive tool for neuromodulation in clinical contexts. Different contexts might call for different patterns of activity modulation. For example, following stroke, alternative therapeutic strategies could involve up or down-regulation of activity in the ipsilateral motor cortex. However, effects of such strategies on activity and brain structure are unknown. In a proof of concept study in healthy individuals, we showed that fMRI neurofeedback can be used to drive activity up or down in ipsilateral motor cortex during hand movement. Given evidence for activity-dependent white matter plasticity, we also tested effects of activity modulation on white matter microstructure using diffusion tensor imaging (DTI). We show rapid opposing changes in corpus callosum microstructure that depend on the direction of activity modulation. Bidirectional modulation of ipsilateral motor cortex activity is therefore possible, and results not only in online changes in activity patterns, but also in changes in microstructure detectable 24 hours later.

## Introduction

Many neuropsychiatric and neurological conditions are associated with aberrant patterns of brain activity so neuromodulation approaches that drive activity towards more favourable patterns are of therapeutic interest. Neurofeedback (NF) taps into the brain’s intrinsic capacity for activity modulation and has been used in a variety of clinical conditions (Sitaram et al., 2017; Wang et al., 2018). Functional magnetic resonance imaging (fMRI) NF provides a powerful approach for spatially specific alteration in brain activity patterns and has been successfully used in in humans at rest (Shibata et al., 2011; Ramot et al., 2016), and during executed movements (Neyedli et al., 2018) or cognitive tasks (Young et al., 2017), with studies showing encouraging behavioural or physiological effects (Sitaram et al., 2017).

Following stroke, there are alterations in activity across the motor system, with changes in activity in ipsilateral (contralesional) motor cortex a particular focus of interest (Johansen-Berg et al., 2002a). Whether this activity should be suppressed or amplified is a matter of debate and the optimal approach might well vary between patients, with suppression appropriate in less impaired patients and amplification in more impaired patients. While there are several studies testing bidirectional modulation of the motor system (in both healthy individuals and stroke patients) using non-invasive brain stimulation (Nitsche and Paulus, 2000; Hesse et al., 2007; Stagg et al., 2009; Lindenberg et al., 2010; Allman et al., 2016; Strube et al., 2016), the few NF studies that targeted ipsilateral motor regions activity have thus far focused on suppressing it only (Auer et al., 2015; Neyedli et al., 2017; Wang et al., 2018). Furthermore, the vast majority of previous studies using NF in the motor system (whether in clinical or healthy populations) have used motor imagery (deCharms et al., 2004; Chiew et al., 2012), but motor execution is arguably more relevant to rehabilitation. Therefore, the current study aimed to address the degree to which activity in ipsilateral motor cortex can be bidirectionally modified with fMRI NF in healthy individuals during executed movements as a proof of concept test of how this approach might be applied and tailored to the patient as an adjunct to neurorehabilitation.

Additionally, we explored the effects of NF-driven activity modulation on white matter microstructure. In humans, studies using diffusion tensor imaging (DTI) have shown that both long-term and short-term learning changes the structure of white matter pathways (Scholz et al., 2009; Hofstetter et al., 2013). Most NF studies have focused on behavioural and functional brain effects, and only two, including a EEG-NF study, have assessed NF effects on the structure of long-range connections (Ghaziri et al., 2013; Marins et al., 2019). While DTI-derived measures such as fractional anisotropy (FA) are nonspecific and modulated by a variety of white matter features, changes in white matter with learning and experience have been related in part to myelin increases in rodents (Sampaio-Baptista et al., 2013; Sampaio-Baptista et al., 2020).

There is growing evidence that myelination can be bidirectionally altered by neuronal activity (Demerens et al., 1996; Stevens et al., 1998) suggesting a bidirectionally-sensitive activity-dependent mechanism might be the underlying driver of learning-related white matter changes (Fields, 2005). Bidirectional NF modulation of focal activity, in combination with DTI measures of white matter microstructure, provide a powerful approach to test these effects in humans. We therefore employed real-time NF, using fMRI at 7 Tesla, to manipulate the activity of the sensorimotor cortices (S1M1s) in opposite directions in two separate conditions, and tested for effects on white matter structure against a sham group.

## Results

Each participant was scanned 4 times and experienced two different NF conditions (only one NF condition was experienced in each session), with DTI acquired before each condition and again 24 hours later (Fig. 1A). During NF, participants were instructed to modulate the height of two bars (representing activity in ipsilateral and contralateral S1M1) on a visual display, by moving *only* their left hand during 30s movement blocks, which alternated with 30s rest blocks. In the ‘Association condition’ participants were required to co-activate both S1M1s (Fig. 1B, C), while in the ‘Dissociation condition’ they were required to maximize contralateral S1M1 activity, while minimizing ipsilateral S1M1 activity (Fig. 1B, D). Participants in the Sham group received the same instructions but were shown the NF videos of a matched participant in the real NF group, and experienced the same two conditions (Association and Dissociation). 80 scans were successfully completed, 28 participants were enrolled and complete data sets were obtained in 20 participants. For each feedback condition, participants trained for approximately 20 minutes (in 3 or 4 runs of ~6 minutes). EMG was used to monitor hand movements online and as expected the muscle activity of the moving (left) hand was significantly higher than the non-moving (right) hand and was similar between Real NF and Sham groups (Supplementary Fig. 1). A debriefing questionnaire (Supplementary Table 1) revealed that both groups felt in control of the feedback (Supplementary Table 2).

**Figure 1.**
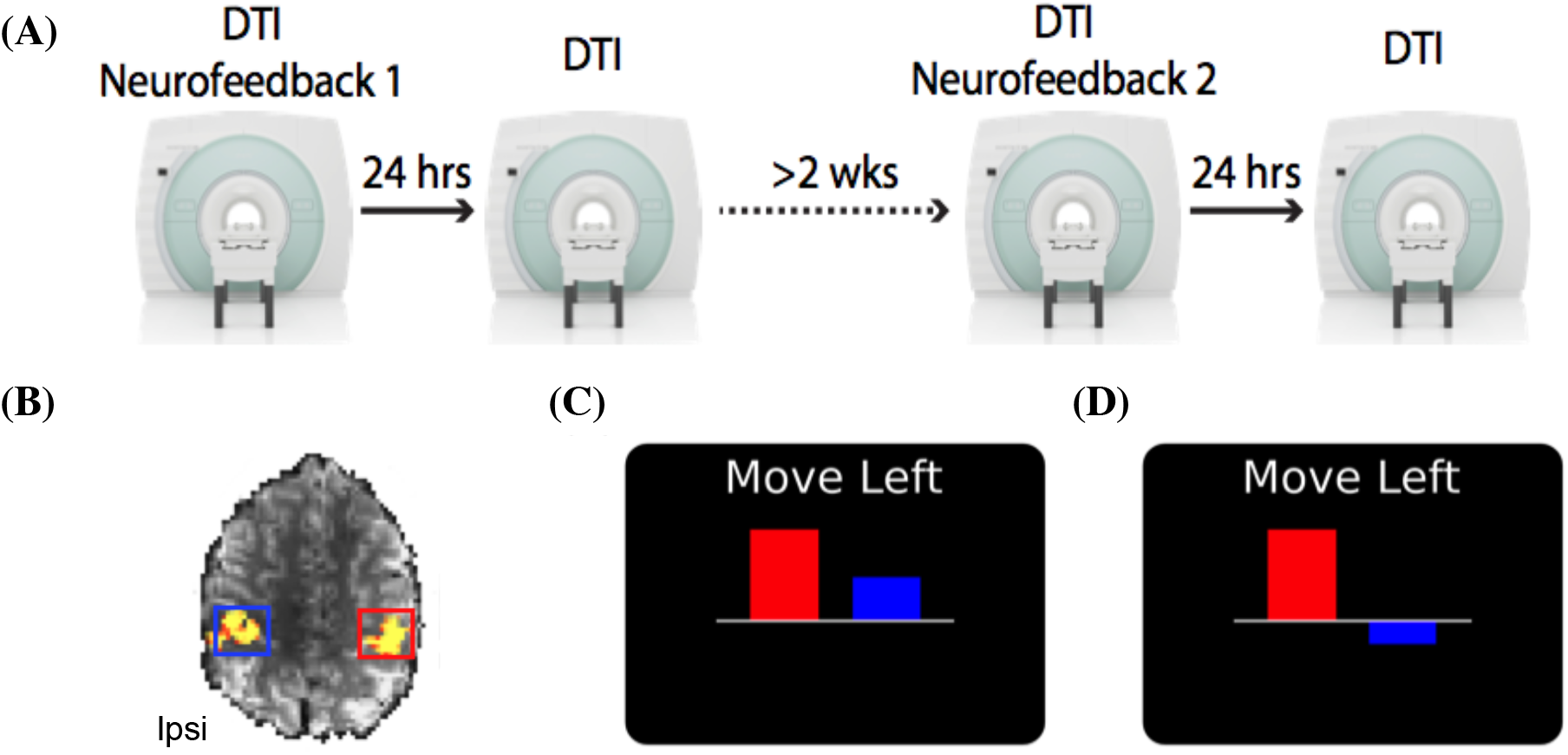
Timeline and neurofeedback (NF) display. (A) Participants in the real NF group and Sham group experienced two different NF conditions, in a counterbalanced design, at least two weeks apart, with DTI acquired before each NF session and again 24 hours later. (B) Functional localizer of an example participant. S1M1 regions of interest were identified by asking the participants to move their right or left fingers sequentially. Ipsi – Ipsilateral (C) Example NF display for the Association condition. (D) Example NF display for the Dissociation condition.

We assessed fMRI activity to test whether participants could modulate S1M1 activity with feedback as instructed. We first analysed signal change within the regions selected during NF using a mixed ANOVA including within-subject factors of condition (Association, Dissociation) and time (Run 1, 2, 3) and between-subject factor of group (Real, Sham). Instructions required participants to increase ipsilateral S1M1 (iS1M1) activity for the Association condition and decrease it for the Dissociation condition. Compared to the Sham group, participants in the NF group were able to modulate activity in iS1M1 as instructed (Fig. 2A, B; main effect of condition: F_(1,18)_ = 8.53, p = 0.009; condition x group interaction: F_(1,18)_ = 12.082, p = 0.003; condition x time x group interaction F_(2,36)_ = 5.03, p = 0.012). Additionally, participants within the NF group were able to modulate iS1M1 activity in opposite directions, with greater, and increasing, activity in the Association condition compared to the Dissociation condition (Fig. 2A: main effect of condition: F_(1,9)_ = 32.045, p = 0.00031; condition x time interaction: F_(2,18)_ = 6.665, p = 0.007). By contrast, no effects of (or interactions with) group were found for the contralateral S1M1 region of interest (ROI), with both groups strongly activating this ROI for both conditions (Fig. 2D,E). Voxel-wise analysis within the NF group revealed specific clusters of significantly greater activity in motor areas, including the ipsilateral hand knob, in the Association compared to the Dissociation condition (Fig. 2C). No other significant clusters were found. The fMRI results therefore demonstrate that the two NF conditions differ in iS1M1 activity, with greater activity seen in the Association condition compared to the Dissociation condition. No significant clusters were found in the sham group for the same comparison, showing that sham participants did not differentially modulate their brain activity between conditions.

**Figure 2.**
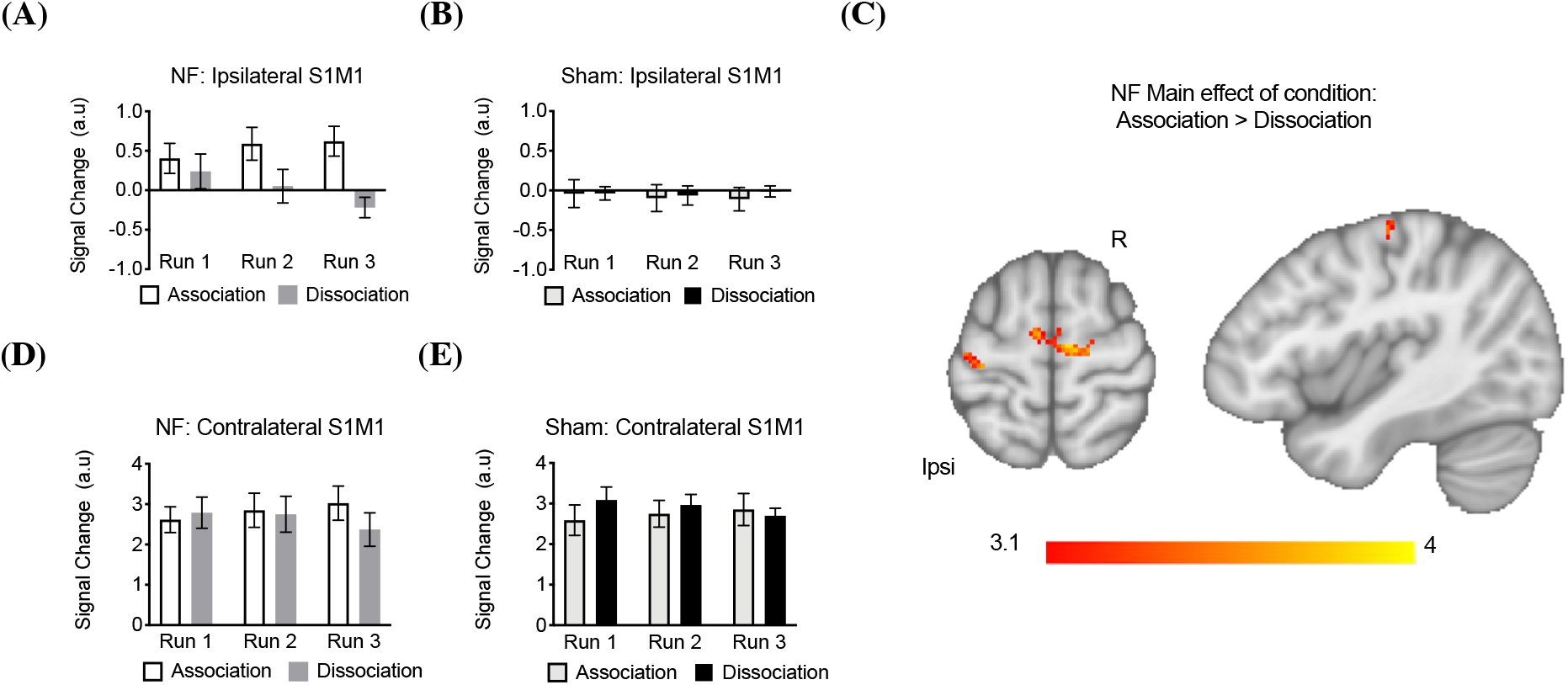
Participants were able to modulate iSIMlactivity with feedback. (A) Participants in the NF group had lower iS1M1 activity in the Dissociation condition compared to the Association condition (B) iS1M1 activity of the Sham group over the 3 runs (C) Voxel-wise analysis showing significantly higher activity in the Association condition compared to the Dissociation condition in iS1M1 (p < 0.05, corrected). (D-E) Instructions required participants to increase contralateral S1M1 (cS1M1) activity for both conditions and both groups. Results showed no main effects or interactions with group, and no main effects of time or condition on signal change within the cS1M1 ROI. There was a significant interaction effect of condition x time which was further explored (F_(2,36)_ = 6.25, p = 0.005). This was driven by an effect of time for the Dissociation condition (F_(2,36)_ = 5.309, p = 0.01), showing both groups decrease activity over time within this condition. No effects of time were identified for the Association condition (F_(2,36)_ = 2.115, p = 0.135). (D) Neurofeedback group contralateral activity in the S1M1 ROI over the 3 runs. (E) Sham group contralateral activity in the S1M1 ROI over the 3 runs. A.u. – Arbitrary units. Ipsi – Ipsilateral Hemisphere, R - Right. Error bars represent SEM.

We went on to test whether this activity modulation resulted in alterations in white matter, as measured by FA, an indirect measure of white matter microstructure previously shown to be sensitive to learning-related white matter plasticity (Scholz et al., 2009; Sampaio-Baptista et al., 2013). Voxel-wise FA maps were calculated from DTI scans acquired before and 24 hours after each NF condition. To accommodate the mixed design nature of this study, FA change maps (post-pre each condition) were first calculated for each condition and each group. Then maps of the difference in FA change between conditions (Dissociation condition FA change – Association condition FA change = condition difference) were calculated for each subject. The resulting maps were then compared between groups.

We used a data-driven approach and performed whole-skeleton voxel-wise non-parametric permutation testing of these between-group differences, which revealed a statistically significant cluster in the corpus callosum, no other clusters were identified elsewhere in the brain (Fig. 3A) (p < 0.05, corrected). This was driven by greater differences between conditions in FA change in the NF group compared to the Sham group and reflected a positive FA change for the Association condition and a negative FA change for the Dissociation condition within the NF group (Fig. 3A). Tractography from this cluster identified pathways connecting sensorimotor and parietal cortices (Supplementary Fig. 2).

**Figure 3.**
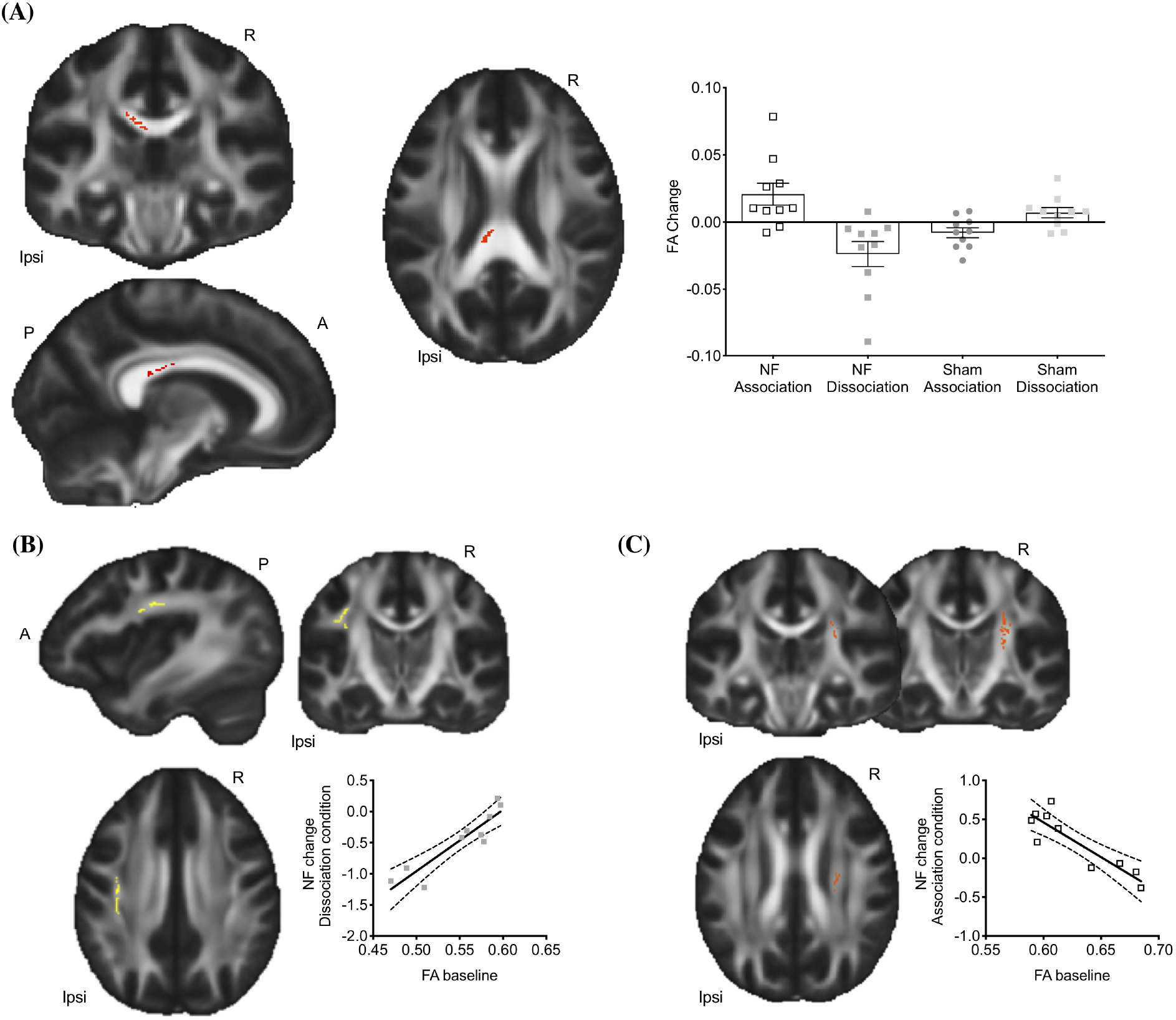
NF training resulted in changes in white matter FA in the corpus callosum. (A) Significant FA cluster (in red) of the between-group contrast (p = 0.05, corrected). Plot on the right side represents the individual participant mean FA change values within significant cluster represented in the FA map and is shown for visualization of range of values and effect direction and not for inference. (B) Significant positive correlation (p < 0.05, corrected) between baseline FA and NF fMRI change for the Dissociation condition in the ipsilateral (left) superior longitudinal fasciculus (SLF) (represented in yellow). (C) A trend (represented in red) towards a significant negative correlation (p = 0.06, corrected) between baseline FA and NF change for the association condition was found in the contralateral (right) corticospinal tract. Plots in (B) and (C) are shown for visualization of range of values and not for inference. Significant cluster are superimposed on the FMRIB FA template. Ipsi – Ipsilateral Hemisphere, A-Anterior, P-Posterior, R - Right. Error bars represent SEM.

We expected that changes in white matter structure would reflect successful modulation of activity with neurofeedback. For each participant we therefore identified which of the two neurofeedback conditions they performed best (see Supplementary Table 3 for more details). We found a significant correlation between the fMRI activity change (run 3 – run 1) and change in FA following this training session (Post24hrs – Baseline) (r = 0.72, p = 0.02, 2-tail). No such correlation was found for the worse neurofeedback session (r = – 0.16, p = 0.66, 2-tail). These correlations were significantly different from each other (test of the difference between two dependent correlations; z = 2.058, p = 0.039, 2-tail). This shows that, following effective neurofeedback training, changes in structure are related to how effectively the participant modulated iS1M1 activity.

Additionally, within the real NF group we tested whether baseline measures of FA correlated with change in fMRI (run 3 – run 1) for each condition. Voxel-wise analysis of the whole skeleton revealed a significant positive correlation (p < 0.05, corrected) between baseline FA and NF fMRI change for the Dissociation condition in the ipsilateral (left) superior longitudinal fasciculus (SLF) (Fig. 3B), suggesting that participants with higher baseline FA in this tract were less able to use neurofeedback to reduce ipsilateral (left) sensorimotor activity as instructed. A trend towards a significant negative correlation (p = 0.06, corrected) between baseline FA and NF change was found in the contralateral (right) corticospinal tract for the Association condition (Fig. 3C), suggesting that higher corticospinal FA at baseline is associated with lower performance when instructed to increase ipsilateral activity with neurofeedback.

## Discussion

Our results support the hypothesis that bidirectional activity modulation of ipsilateral sensorimotor activity during executed hand movement can be achieved via neurofeedback and that this results in rapid, directional, and anatomically specific changes in white matter structure. This finding in healthy individuals is relevant to considering application of neurofeedback in therapeutic contexts and in particular as an adjunct to motor neurorehabilitation after stroke.

The two conditions here could potentially be used as alternative interventions in stroke patients. For instance, rebalancing of aberrant cortical activity by decreasing motor activity of the ipsilateral (spared) hemisphere is a potential route for improving motor function after stroke particularly in patients with low levels of impairment (Johansen-Berg et al., 2002b; Johansen-Berg, 2003; Johansen-Berg, 2007), while enhancing activity in the ipsilateral motor cortex may be beneficial for patients with more severe impairment (Bradnam et al., 2012; McDonnell and Stinear, 2017). Future studies should assess these approaches in patients, where behavioural, motor evoked-responses (MEPs) and MRI markers (Stinear et al., 2007) could be used as predictors for tailoring the NF intervention to the individual, including which brain areas to target and in which direction.

Activity in the ipsilesional S1M1 ROI showed clear modulation with NF, while activity in the contralateral ROI remained fairly similar across NF and sham conditions, despite both real NF conditions instructing increases in contralateral activity. Given that hand movements strongly elicit contralateral sensorimotor activity in healthy participants it is likely that this is due to a ceiling effect, with gains in activity not easily achieved. By contrast, there is typically very little activity in ipsilateral S1M1 during this task in healthy individuals and so more scope for modulation. Future studies employing several training sessions should assess whether further training leads to progressive increases in contralateral activity, particularly in patient groups with motor deficits or in older populations who may have lower activity in this region at baseline.

Our white matter results suggest that the focal activity modulation evoked by NF resulted in changes in white matter microstructure that could be detected 24 hours later. Likely more than one cellular mechanism underlies our structural findings. FA is modulated by several white matter features such as myelination, axon density and caliber, and potentially by astrocyte morphology, cell swelling, or changes in membrane fluidity (Sampaio-Baptista and Johansen-Berg, 2017). Myelination is an attractive potential mechanism because neuronal activity can bidirectionally regulate myelin formation and compaction and OPCs proliferation and differentiation (Demerens et al., 1996; Stevens et al., 1998; Gibson et al., 2014) and such effects occur over similar timescales to the ones used here (Xiao et al., 2016). For example, 30 minutes of optogenetic stimulation of premotor neurons led to rapid increases in oligodendrocyte precursor cells (OPCs) proliferation and differentiation within 24 hours (Gibson et al., 2014). Furthermore, recently matured oligodendrocytes can form, extend and retract myelin segments within 24 hours (Watkins et al., 2008; Czopka et al., 2013). Preexisting oligodendrocytes could also contribute to myelin remodeling (Yeung et al., 2014; Dutta et al., 2018; Mitew et al., 2018), however strong evidence for this process is currently scarce and the timescale at which this occurs is unknown.

Importantly, new oligodendrocytes, formed during adulthood, play an essential role in new motor skill acquisition (McKenzie et al., 2014), with impairments detected within a couple of hours (Xiao et al., 2016), suggesting that learning is supported not only by neuronal changes, such as synaptic plasticity, but also by adjunct changes in myelination (Long and Corfas, 2014). Concurrently, other white matter structural properties such as axon density and caliber, astrocyte morphology, cell swelling, or changes in membrane fluidity also occur in response to neuronal activity and learning and could underlie some of the DTI effects (Blumenfeld-Katzir et al., 2011; Sampaio-Baptista and Johansen-Berg, 2017; Sinclair et al., 2017).

The structural findings suggest that alterations in the elicited brain activity is a possible mediator of previously described experience-related white matter changes in the human brain resulting from behavioural interventions (Scholz et al., 2009; Hofstetter et al., 2013). However, given the complex nature of the BOLD signal it is not straightforward to attribute BOLD fMRI increases or decreases to either net excitation or net inhibition (Devor et al., 2007; Logothetis, 2008), though some cellular specific mechanisms that contribute to BOLD have been recently described (Uhlirova et al., 2016).

The anatomical site of the detected FA changes indicates that successful modulation of left sensorimotor activity by performing left hand movements resulted in changes mainly in the fibres that connect to the opposite hemisphere. This suggests that modulation of ipsilateral activity occurred via callosal connections, resulting in structural alterations in the connections between the two cortices. The corpus callosum is a relatively coherent fibre bundle as such small changes might be easier to detect in this location, whereas white matter closer to the cortex contains more crossing fibres and so effects of structural modulation on DTI metrics could be harder to detect.

One challenge with clinical application of NF is that there is high variability in neurofeedback success. Many studies identify ‘responders’ and ‘non-responders’ to NF and the individual factors that determine NF success are not well understood. Here, we tested whether any baseline variables correlated with neurofeedback success. We found that in the Dissociation condition, higher ipsilateral SLF FA at baseline is associated with higher ipsilateral sensorimotor fMRI change, while higher contralateral corticospinal FA at baseline is associated with lower performance in increasing ipsilateral activity with neurofeedback. Both these results indicate that high FA in motor-related pathways is associated with worse neurofeedback performance, implying that highly structurally connected motor networks might be harder to modulate via neurofeedback. These results open the possibility of using structural imaging to predict neurofeedback performance, explain inter-individual variability and potentially tailor the intervention to the needs of the participants. For instance, more neurofeedback sessions might be necessary to change the activity of a highly structurally connected network.

## Methods

### Participants and design

All participants provided written informed consent in accordance with the University of Oxford ethics committee approval of the protocol (MSD-IDREC-C1-2012-151). 28 right-handed participants (22-38 year old, 15 female) were recruited and scanned in a 7T Siemens scanner 4 times (Fig. 1A).

Unbeknownst to the participants they were assigned to two different groups: Neurofeedback or Sham. Total number of analyzed scans was 80 (NF group n=10×4=40; sham group n=10×4=40). One participant in the NF group did not complete the experiment due to back pain and DTI was not acquired in 1 participant due to scanner crashes. Four participants in the Sham group did not complete the full study and data was not fully collected in further 2 Sham participants due to scanner crashes.

Participants were blind to group assignment. The experimenter could not be blinded to group due to limitations of the real time software. However, all participants received identical instructions and experimental procedures were the same across the two groups with the exception of the source of the feedback presented (see below). Each participant in each group was scanned under two experimental conditions: association and Dissociation. For each condition participants were scanned twice, 24 hours apart. The order of experimental conditions was counterbalanced across participants and conditions were spaced at least 2 weeks apart (mean = 33.8 days, SD=16.3) (see Fig. 1A).

### MRI data acquisition

Imaging was performed on a 7.0T Siemens Magnetom MRI system (Siemens, Erlangen, Germany) with a 32-channel head coil at the FMRIB Centre at the University of Oxford. For each condition, scans were acquired over two days as follows:

#### Day 1: FMRI Neurofeedback and DTI

All FMRI data was acquired with a gradient echo planar image sequence (16 slices, 2 mm axial plane, 2 x 2 mm^2^ in plane resolution, repetition time (TR)=2000 ms; echo time (TE)=25 ms; flip angle=90°). A whole brain echo planar image sequence was acquired for registration purposes (60 slices, 2 mm axial plane, 2 x 2 mm^2^ in plane resolution, TR=3500 ms; echo time=25 ms; flip angle=90° field of view, 220 x 220mm).

#### Functional localizer

The functional localizer consisted of eight, 12-second tapping blocks, four blocks for each hand, interspersed with 24 seconds rest. The participants saw the instructions ‘Right Tap’, ‘Left Tap’ and ‘Rest’ displayed in white on a black background. Participants were told to use each finger in sequence starting with their index finger and moving outward towards the little finger at a rate of approximately 1 Hz, and to repeat the sequence until they saw the rest instruction. The results from the real-time general linear model (GLM) of the localizer scan were used to select two motor ROIs (18 x 18 x 10mm) for each participant, each centered over the peak of activation in both hemispheres.

#### DTI

After the functional localizer we acquired whole brain diffusion-weighted volumes (64 directions; b-value = 1500 s/mm^2^; 80 slices; voxel size 1.5 x 1.5 x 1.5 mm^3^; TR=10 s; TE= 64 ms) including 5 volumes without diffusion weighting (b-value = 0 s/mm^2^)), and also a separate data set without diffusion weighting with opposite phase encoding for correction of susceptibility induced distortions (b-value = 0 s/mm^2^, 80 slices; voxel size 1.5 x 1.5 x 1.5 mm^3^; repetition time (TR) = 10 s; echo time (TE) = 64 ms). The total acquisition time for DTI was about 10 minutes.

#### Neurofeedback training

Next, three, ~6 min feedback (FB) functional scans were acquired with a block design (30 seconds on, 30 seconds rest). In five sessions a fourth FB scan was acquired but only the first 3 training runs that were common to all participants were analysed.

Turbo-BrainVoyager software version 3.2 (Brain Innovation, Maastricht, The Netherlands; Goebel, 2001) was used to calculate the BOLD signal in real time using a whole-brain voxelwise recursive GLM. The feedback signal was based on the averaged voxel time-course extracted from the localized motor ROIs and the feedback image was updated each TR (2 seconds). Online motion correction in three dimensions, including translations and rotations were used to correct for head movements during the scan.

A custom-made transmission control protocol (TCP) based network interface plug-in for Turbo-Brain Voyager was used to transmit the preprocessed ROI time-course to Turbo-Feedback, a custom-made software tool, which performed neurofeedback signal calculation and displayed the resulting feedback to the participants.

Participants saw two vertical bars, representing the contralateral and ipsilateral hemispheric activity, with a horizontal line delineating the center point. The equation used to calculate the height of each bar was the following:

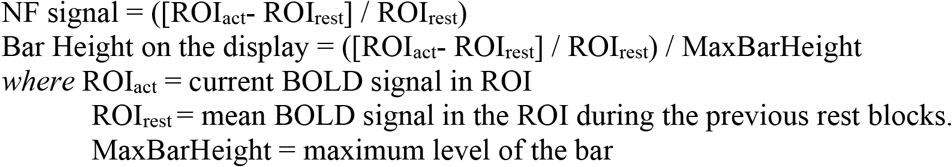

Positive values were represented above the centre point, and negative values below the centre point (Fig 1C,D). The bar for contralateral (right hemisphere) activation was displayed on the left as this hemisphere should be most active during left hand movement. The bar for ipsilateral (left hemisphere) activation was displayed on the right.

#### Sham Group

Participants in the Sham group were matched to a participant in the NF group and received feedback videos from that participant (rather than their feedback from their own brain activity). This allowed the Sham participants to have a similar experience as the NF group. All scans and instructions received by the Sham group were identical to those received by the NF group.

#### Experimental conditions

Two neurofeedback conditions were tested in separate sessions in both the NF and the Sham group:

*Association condition* Participants were instructed to increase the size of both bars *Dissociation condition* Participants were instructed to decrease the size of the bar on the right side of the screen, while increasing the bar size on the left side

In this way, the goal of the Association condition was to maximise activation in both left and right S1M1, whereas the goal of the Dissociation condition was to maximise right (contralateral) S1M1 activity and minimize left (ipsilateral) S1M1 activity.

Participants were only told that the bars represented their brain activity. For both conditions participants were asked to perform left hand movements in order to modulate the height of the bars. The participants saw the instructions ‘Move Left’ or ‘Rest’ displayed in white on a black background. Participants were allowed to use any left hand movement strategy to accomplish the goal in each condition. Participants were instructed not to move their right hand during feedback training and both arms were monitored on-line for movement using EMG. During the instructions a list of example strategies were read but participants were told they could use any other strategy as long as they did not move their right hand:

> *“Open and close hand, move fingers, make grasping movements, move fingers sequentially or randomly, imagine hand/finger movements, focus on moving hand or non-moving hand, increase rate, force, size, of movement etc.”*

#### Day 2: DTI

24 hours after each NF training session DTI was again acquired with the same parameters as above. For registration purposes, one structural image per subject was acquired during the second session only using a T1 weighted, MPRAGE sequence with 1 x 1 x 1 mm^3^ isotropic voxels (TR=2200 ms; TE=2.2 ms; flip angle 7°, field of view, 192×192; matrix=192×192).

### EMG Acquisition

A Biopac system and AcqKnowledge software (Version 4.2) were used for EMG acquisition during neurofeedback sessions. Due to technical difficulties we only acquired a full set of EMG data in 7 participants in the NF group and 9 participants in the Sham group. We used two MRI safe surface electrodes (ConMed corporation, USA) to record from the flexor carpi ulnaris muscle and an additional electrode placed over the elbow olecranon was used as the ground electrode. AcqKnowledge software was used to monitor and record muscle activity during the feedback training acquisition with online MRI artifact and line noise correction.

### Neurofeedback questionnaire

Following each feedback training session participants completed a questionnaire outside the scanner (Supplementary Table 1). Participants reported on a scale of 1-5 how much control they felt they had over the bar. Then a number of strategies for controlling the FB were presented and participants were asked to report if they used the strategy and, if so, how successful they thought the strategy was on a scale of 1-5.

## Data analysis

### fMRI Preprocessing

BOLD fMRI data for each subject were analyzed with FMRIB’s expert analysis tool (FEAT, version 5.98) from the FMRIB software library version 5.0 (www.fmrib.ox.ac.uk/fsl). Preprocessing of the images included motion correction with FMRIB’s Linear Image Registration Tool (MCFLRIT), brain extraction with BET, spatial smoothing using a Gaussian kernel of 5 mm FWHM, and highpass temporal filtering of 150 s.

Functional data were first aligned to the whole brain scan and then to the subject’s structural image with linear registration (FMRIB’s Linear Image Registration Tool, FLIRT), and then optimized using Boundary-Based Registration (Greve and Fischl, 2009). For structural images we used the anatomical processing script (fsl_anat) to robustly correct the bias-field and register the images to standard MNI space. The resulting warp fields were then applied to the functional images.

We used a voxel-based general linear model (GLM), as implemented in FEAT. For each neurofeedback training scan, the block design paradigm (30 s hand movement plus feedback and 30 s rest) convolved with a gamma function, along with its temporal derivative, was used to model the activation time course.

### ROI fMRI analysis of the Feedback Training

After first-level Feat analysis, the tool featquery was used to extract the percentage signal change of the defined motor ROIs. Mixed design ANOVA or Repeated-Measures ANOVA (SPSS version 25) were used when appropriate to test for main effects of group, condition (Association, Dissociation), NF run (1, 2, 3) and interaction effects between these variables. The significance threshold used was p < 0.05.

### Group-level voxel-wise fMRI analysis

To test for main effects of condition we used a within-subject fixed-effects second-level analysis to calculate the average activation for the contrast of movement versus rest across the three feedback scans per participant. The resulting maps were then fed into group level analysis using FMRIB’s Local Analysis of Mixed Effects (Woolrich et al., 2004).

We tested for differences between Association and Dissociation conditions with a paired t-test. Z statistic images were thresholded using clusters determined by Z > 3.1 and a family-wiseerror-corrected cluster significance threshold of p < 0.05 was applied to the suprathreshold clusters.

### DTI analysis

DTI data were analysed with FMRIB’s Diffusion Toolbox (FDT). Two sets of volumes without diffusion-weighting were collected, with reversed phase-encode blips (i.e., one set with anterior-posterior encoding and one with posterior-anterior), resulting in pairs of images with distortions going in opposite directions. From these image pairs the susceptibility-induced off-resonance field was estimated using a method similar to that described in (Andersson et al., 2003) as implemented in FSL (Smith et al., 2004) All data were then corrected for susceptibility induced distortions and for eddy current distortions and head movements with the FSL’s eddy tool. A diffusion tensor model was then fit to the data at each voxel using dtifit and voxel-wise maps of fractional anisotropy (FA), mean diffusivity (MD), radial and axial diffusivity were estimated for each subject and each timepoint. These maps were then analysed using Tract Based Spatial Statistics (TBSS) (Smith et al., 2006). We performed unbiased registration by registering the maps to the study specific template.

A mixed-design ANOVA is not accommodated by the general linear model (GLM) as implemented in the FSL tool Randomise (http://fsl.fmrib.ox.ac.uk/fsl/fslwiki/Randomise). As such to be able to test for group differences we have first computed the FA change (post-pre) maps for each condition and each participant. We then calculated the difference between conditions for each participant (Dissociation FA change – Association FA change = condition difference). These maps were then compared between groups (NF vs Sham) with an unpaired t-test. This allows us to test whether differences in FA change between conditions were greater in magnitude in the NF group compared to the Sham group. Gender was used as a covariate (2/8 males/females in NF group and 4/6 males/females in the Sham group).

We tested for group differences with an unpaired t-test by feeding these difference maps into Randomise for permutation-based non-parametric testing of whole-skeleton FA. Clusters were formed at t > 1.7 and tested for significance at p < 0.05, corrected for multiple comparisons across space (Nichols and Holmes, 2002).

### Correlations between fMRI change and FA change

We wished to test whether subjects who showed the most effective FMRI modulation with feedback also had the greatest microstructural change in white matter. To do so, we first calculated change in the fMRI activity (Run 3 – Run 1) for the iS1M1 ROI for each condition. Rather than consider both conditions for each participant, we selected for each participant the condition in which they performed best. By considering only one condition per participant we could also ensure independence of data points for correlation calculation. Best performance was defined as highest activity change in the instructed direction. 40% of the participants responded best to the Association condition and 60% to the Dissociation condition, 50% of the participants performed best in the first session regardless of condition (Supplementary Table 3). We tested for correlations between this iS1M1 fMRI change and the corresponding FA change (Post24hrs-Baseline) with Spearman’s Rho (p < 0.05, 2-tail) (SPSS version 25).

### Tractography analysis

We used tractography to identify the probabilistic connectivity map of the significant corpus callosum FA cluster (i.e. cluster shown in Fig. 3A). First, for each participant, BEDPOSTX was used to automatically determine the number of estimated fiber populations per brain voxel and to fit estimates of principle diffusion direction for each population (Behrens et al., 2007). Then PROBTRACKX (5000 samples, 0.5 mm step length, 2000 steps, 0.2 curvature threshold) was used to follow these estimates in order to generate a probabilistic connectivity distribution, using the FA cluster as a seed. The resulting individual participant probabilistic connectivity maps were thresholded at 100. We then created two maps to illustrate the connectivity of the significant FA cluster. To create the mean probability map the individual maps were then overlapped across participants and the mean was extracted (Supplementary Fig. 2A). To represent the tracts common to the population, the maps were binarized, added together and colour-coded (Supplementary Fig. 2B).

### EMG analysis

EMG data were band pass filtered offline from 20 Hz to 200 Hz, full-wave rectified and converted to root mean square (RMS) using a 50 ms window period. For statistical comparison, response-locked RMS-EMG activity was averaged from 0 to 30 s for each movement block, after subtracting the 3 s before each movement onset as baseline. We used a Mixed Design ANOVA (SPSS version 25) to test for effects of group, condition, run and hand.

### Questionnaire analysis

A Wilcoxon signed-rank test was conducted to compare how much control the participants felt they had over the feedback between conditions within group (Question A, Supplementary Table 1). A Mann-Whitney U test was used to compare how much control the participants felt they had over the feedback between groups (Question A, Supplementary Table 1). A Mann-Whitney U test was used to test if there were differences between groups in how successful the strategies were perceived to be (Question B, Supplementary Table 1).

## Acknowledgments

This work was supported by the Wellcome Trust (WT090955AIA and 110027/Z/15/Z to H J-B). The research was further supported by Marie Curie Actions (Adaptive Brain Computations network PITN-GA-2008-290011) and by the National Institute for Health Research (NIHR) Oxford Biomedical Research Centre based at Oxford University Hospitals NHS Trust and University of Oxford. The views expressed are those of the author(s) and not necessarily those of the NHS, the NIHR or the Department of Health. The Wellcome Centre for Integrative Neuroimaging is supported by core funding from the Wellcome Trust (203139/Z/16/Z). The authors would like to thank N. Filippini, S. Foxley and the radiography team for operating the 7T scanner, T. Makin and F. Eippert for assistance with the EMG, S. Sotiropoulos for assistance with the 3D tractography pictures, and W. Richardson, T. Makin and J. O’Shea for the useful comments on the manuscript.

## Author contribution

C.S-B. designed the study, collected and analysed the data. H.N. collected data and provided assistance with data analysis. Z.B.S. collected and analysed data. D.H and K. D. collected data. Y.H. provided assistance with EMG data analysis. J.A., M.L. and R.G. developed and provided assistance with data collection methods and analysis. H.J-B designed the study and supervised the project. C.S-B wrote the manuscript and all authors edited it.

## References

Allman C, Amadi U, Winkler AM, Wilkins L, Filippini N, Kischka U, Stagg CJ, Johansen-Berg H (2016) Ipsilesional anodal tDCS enhances the functional benefits of rehabilitation in patients after stroke. Sci Transl Med 8:330re331.

Andersson JL, Skare S, Ashburner J (2003) How to correct susceptibility distortions in spin-echo echo-planar images: application to diffusion tensor imaging. Neuroimage 20:870–888.

Auer T, Schweizer R, Frahm J (2015) Training Efficiency and Transfer Success in an Extended Real-Time Functional MRI Neurofeedback Training of the Somatomotor Cortex of Healthy Subjects. Front Hum Neurosci 9:547.

Behrens TEJ, Berg HJ, Jbabdi S, Rushworth MFS, Woolrich MW (2007) Probabilistic diffusion tractography with multiple fibre orientations: What can we gain? Neuroimage 34:144–155.

Blumenfeld-Katzir T, Pasternak O, Dagan M, Assaf Y (2011) Diffusion MRI of structural brain plasticity induced by a learning and memory task. PLoS One 6:e20678.

Bradnam LV, Stinear CM, Barber PA, Byblow WD (2012) Contralesional hemisphere control of the proximal paretic upper limb following stroke. Cereb Cortex 22:2662–2671.

Chiew M, LaConte SM, Graham SJ (2012) Investigation of fMRI neurofeedback of differential primary motor cortex activity using kinesthetic motor imagery. Neuroimage 61:21–31.

Czopka T, Ffrench-Constant C, Lyons DA (2013) Individual oligodendrocytes have only a few hours in which to generate new myelin sheaths in vivo. Dev Cell 25:599–609.

deCharms RC, Christoff K, Glover GH, Pauly JM, Whitfield S, Gabrieli JD (2004) Learned regulation of spatially localized brain activation using real-time fMRI. Neuroimage 21:436–443.

Demerens C, Stankoff B, Logak M, Anglade P, Allinquant B, Couraud F, Zalc B, Lubetzki C (1996) Induction of myelination in the central nervous system by electrical activity. Proc Natl Acad Sci U S A 93:9887–9892.

Devor A, Tian P, Nishimura N, Teng IC, Hillman EM, Narayanan SN, Ulbert I, Boas DA, Kleinfeld D, Dale AM (2007) Suppressed neuronal activity and concurrent arteriolar vasoconstriction may explain negative blood oxygenation level-dependent signal. J Neurosci 27:4452–4459.

Dutta DJ, Woo DH, Lee PR, Pajevic S, Bukalo O, Huffman WC, Wake H, Basser PJ, SheikhBahaei S, Lazarevic V, Smith JC, Fields RD (2018) Regulation of myelin structure and conduction velocity by perinodal astrocytes. Proc Natl Acad Sci U S A 115:11832–11837.

Fields RD (2005) Myelination: an overlooked mechanism of synaptic plasticity? Neuroscientist 11:528–531.

Ghaziri J, Tucholka A, Larue V, Blanchette-Sylvestre M, Reyburn G, Gilbert G, Levesque J, Beauregard M (2013) Neurofeedback training induces changes in white and gray matter. Clin EEG Neurosci 44:265–272.

Gibson EM, Purger D, Mount CW, Goldstein AK, Lin GL, Wood LS, Inema I, Miller SE, Bieri G, Zuchero JB, Barres BA, Woo PJ, Vogel H, Monje M (2014) Neuronal activity promotes oligodendrogenesis and adaptive myelination in the mammalian brain. Science 344:1252304.

Greve DN, Fischl B (2009) Accurate and robust brain image alignment using boundary-based registration. Neuroimage 48:63–72.

Hesse S, Werner C, Schonhardt EM, Bardeleben A, Jenrich W, Kirker SG (2007) Combined transcranial direct current stimulation and robot-assisted arm training in subacute stroke patients: a pilot study. Restor Neurol Neurosci 25:9–15.

Hofstetter S, Tavor I, Tzur Moryosef S, Assaf Y (2013) Short-term learning induces white matter plasticity in the fornix. J Neurosci 33:12844–12850.

Johansen-Berg H (2003) Motor physiology: a brain of two halves. Current Biology 13:R802–R804.

Johansen-Berg H (2007) Functional imaging of stroke recovery: what have we learnt and where do we go from here? Int J Stroke 2:7–16.

Johansen-Berg H, Rushworth MF, Bogdanovic MD, Kischka U, Wimalaratna S, Matthews PM (2002a) The role of ipsilateral premotor cortex in hand movement after stroke. Proc Natl Acad Sci U S A 99:14518–14523.

Johansen-Berg H, Dawes H, Guy C, Smith S, Wade D, Matthews P (2002b) Correlation between motor improvements and altered fMRI activity after rehabilitative therapy. Brain 125:2731.

Lindenberg R, Renga V, Zhu LL, Nair D, Schlaug G (2010) Bihemispheric brain stimulation facilitates motor recovery in chronic stroke patients. Neurology 75:2176–2184.

Logothetis NK (2008) What we can do and what we cannot do with fMRI. Nature 453:869–878.

Long P, Corfas G (2014) Neuroscience. To learn is to myelinate. Science 346:298–299.

Marins T, Rodrigues EC, Bortolini T, Melo B, Moll J, Tovar-Moll F (2019) Structural and functional connectivity changes in response to short-term neurofeedback training with motor imagery. Neuroimage 194:283–290.

McDonnell MN, Stinear CM (2017) TMS measures of motor cortex function after stroke: A meta-analysis. Brain Stimul 10:721–734.

McKenzie IA, Ohayon D, Li H, de Faria JP, Emery B, Tohyama K, Richardson WD (2014) Motor skill learning requires active central myelination. Science 346:318–322.

Mitew S, Gobius I, Fenlon LR, McDougall SJ, Hawkes D, Xing YL, Bujalka H, Gundlach AL, Richards LJ, Kilpatrick TJ, Merson TD, Emery B (2018) Pharmacogenetic stimulation of neuronal activity increases myelination in an axon-specific manner. Nat Commun 9:306.

Neyedli HF, Sampaio-Baptista C, Kirkman MA, Havard D, Luhrs M, Ramsden K, Flitney DD, Clare S, Goebel R, Johansen-Berg H (2017) Increasing lateralized motor activity in younger and older adults using Real-time fMRI during executed movements. Neuroscience.

Neyedli HF, Sampaio-Baptista C, Kirkman MA, Havard D, Luhrs M, Ramsden K, Flitney DD, Clare S, Goebel R, Johansen-Berg H (2018) Increasing Lateralized Motor Activity in Younger and Older Adults using Real-time fMRI during Executed Movements. Neuroscience 378:165–174.

Nichols TE, Holmes AP (2002) Nonparametric permutation tests for functional neuroimaging: a primer with examples. Hum Brain Mapp 15:1–25.

Nitsche MA, Paulus W (2000) Excitability changes induced in the human motor cortex by weak transcranial direct current stimulation. J Physiol 527 Pt 3:633–639.

Ramot M, Grossman S, Friedman D, Malach R (2016) Covert neurofeedback without awareness shapes cortical network spontaneous connectivity. Proc Natl Acad Sci U S A 113:E2413–2420.

Sampaio-Baptista C, Johansen-Berg H (2017) White Matter Plasticity in the Adult Brain. Neuron 96:1239–1251.

Sampaio-Baptista C, Valles A, Khrapitchev AA, Akkermans G, Winkler AM, Foxley S, Sibson NR, Roberts M, Miller K, Diamond ME, Martens GJM, De Weerd P, Johansen-Berg H (2020) White matter structure and myelin-related gene expression alterations with experience in adult rats. Prog Neurobiol:101770.

Sampaio-Baptista C, Khrapitchev AA, Foxley S, Schlagheck T, Scholz J, Jbabdi S, DeLuca GC, Miller KL, Taylor A, Thomas N, Kleim J, Sibson NR, Bannerman D, Johansen-Berg H (2013) Motor skill learning induces changes in white matter microstructure and myelination. J Neurosci 33:19499–19503.

Scholz J, Klein MC, Behrens TE, Johansen-Berg H (2009) Training induces changes in white-matter architecture. Nat Neurosci 12:1370–1371.

Shibata K, Watanabe T, Sasaki Y, Kawato M (2011) Perceptual learning incepted by decoded fMRI neurofeedback without stimulus presentation. Science 334:1413–1415.

Sinclair JL, Fischl MJ, Alexandrova O, Hebeta M, Grothe B, Leibold C, Kopp-Scheinpflug C (2017) Sound-Evoked Activity Influences Myelination of Brainstem Axons in the Trapezoid Body. J Neurosci 37:8239–8255.

Sitaram R, Ros T, Stoeckel L, Haller S, Scharnowski F, Lewis-Peacock J, Weiskopf N, Blefari ML, Rana M, Oblak E, Birbaumer N, Sulzer J (2017) Closed-loop brain training: the science of neurofeedback. Nat Rev Neurosci 18:86–100.

Smith SM, Jenkinson M, Johansen-Berg H, Rueckert D, Nichols TE, Mackay CE, Watkins KE, Ciccarelli O, Cader MZ, Matthews PM, Behrens TE (2006) Tract-based spatial statistics: voxelwise analysis of multi-subject diffusion data. Neuroimage 31:1487–1505.

Smith SM, Jenkinson M, Woolrich MW, Beckmann CF, Behrens TE, Johansen-Berg H, Bannister PR, De Luca M, Drobnjak I, Flitney DE, Niazy RK, Saunders J, Vickers J, Zhang Y, De Stefano N, Brady JM, Matthews PM (2004) Advances in functional and structural MR image analysis and implementation as FSL. Neuroimage 23 Suppl 1:S208–219.

Stagg CJ, Best JG, Stephenson MC, O’Shea J, Wylezinska M, Kincses ZT, Morris PG, Matthews PM, Johansen-Berg H (2009) Polarity-sensitive modulation of cortical neurotransmitters by transcranial stimulation. J Neurosci 29:5202–5206.

Stevens B, Tanner S, Fields RD (1998) Control of myelination by specific patterns of neural impulses. J Neurosci 18:9303–9311.

Stinear CM, Barber PA, Smale PR, Coxon JP, Fleming MK, Byblow WD (2007) Functional potential in chronic stroke patients depends on corticospinal tract integrity. Brain 130:170–180.

Strube W, Bunse T, Nitsche MA, Nikolaeva A, Palm U, Padberg F, Falkai P, Hasan A (2016) Bidirectional variability in motor cortex excitability modulation following 1 mA transcranial direct current stimulation in healthy participants. Physiol Rep 4.

Uhlirova H et al. (2016) Cell type specificity of neurovascular coupling in cerebral cortex. Elife 5.

Wang T, Mantini D, Gillebert CR (2018) The potential of real-time fMRI neurofeedback for stroke rehabilitation: A systematic review. Cortex 107:148–165.

Watkins TA, Emery B, Mulinyawe S, Barres BA (2008) Distinct stages of myelination regulated by gamma-secretase and astrocytes in a rapidly myelinating CNS coculture system. Neuron 60:555–569.

Woolrich MW, Behrens TE, Beckmann CF, Jenkinson M, Smith SM (2004) Multilevel linear modelling for FMRI group analysis using Bayesian inference. Neuroimage 21:1732–1747.

Xiao L, Ohayon D, McKenzie IA, Sinclair-Wilson A, Wright JL, Fudge AD, Emery B, Li H, Richardson WD (2016) Rapid production of new oligodendrocytes is required in the earliest stages of motor-skill learning. Nat Neurosci 19:1210–1217.

Yeung MS, Zdunek S, Bergmann O, Bernard S, Salehpour M, Alkass K, Perl S, Tisdale J, Possnert G, Brundin L, Druid H, Frisen J (2014) Dynamics of oligodendrocyte generation and myelination in the human brain. Cell 159:766–774.

Young KD, Siegle GJ, Zotev V, Phillips R, Misaki M, Yuan H, Drevets WC, Bodurka J (2017) Randomized Clinical Trial of Real-Time fMRI Amygdala Neurofeedback for Major Depressive Disorder: Effects on Symptoms and Autobiographical Memory Recall. Am J Psychiatry 174:748–755.

