## Supplementary Info for "Neurofeedback fMRI in the motor system elicits bi-directional changes in activity and white-matter structure in the healthy adult human brain"

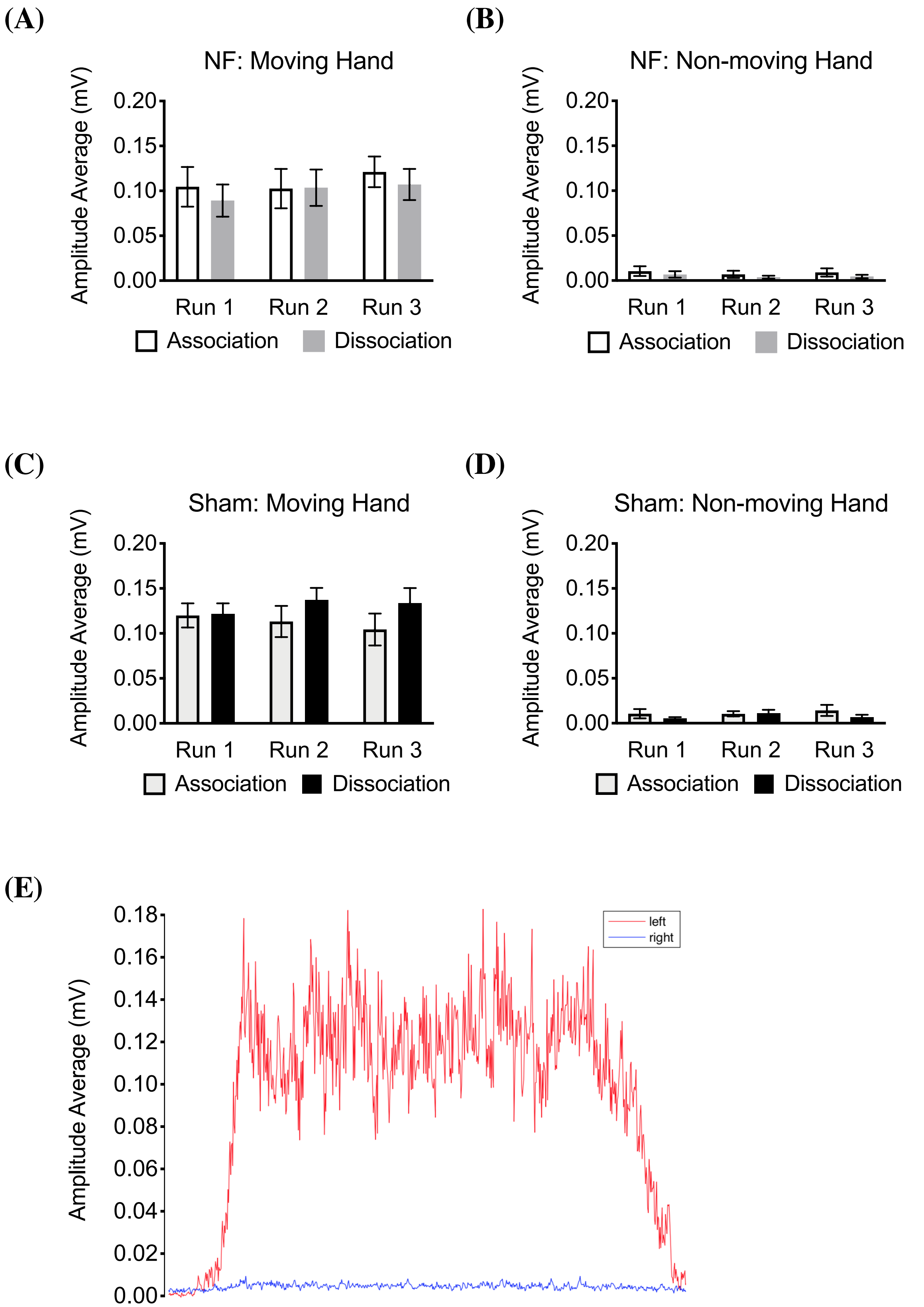


**Supplementary Figure 1. EMG results**. Mixed Design ANOVA that included group, condition, run and hand as factors revealed an effect of hand, with the average amplitude EMG of the moving (left) hand significantly higher than the non-moving (right) hand (Fig S1A, B) ($F_{(1,14)} = 182.472, p < 0.001$). There was no effect of group ($F_{(1,14)} = 0.522$, p =

0.482), condition ($F_{(1,14)} = 0.006$, $p = 0.938$), or run (($F_{(2,28)} = 1.508$, $p = 0.239$) nor any interaction effects (all $p > 0.05$). (A) Average amplitude (mV) of the left hand for the NF group. (B) Average amplitude of the right hand for the NF group. (C) Average amplitude (mV) of the left hand for the Sham group. (D) Average amplitude of the right hand for the Sham group (E) Example from one participant of the average activation for each hand. Error bars represent SEM.

| Questionnaire |
| --- |
| A. How much control over the blue bar did you feel you had? (1-not in control, 5-full in control) |
| B. Which strategies did you use? rate on a scale of 1 to 5 how successful the strategy was (1 – unsuccessful strategy; 5 – successful strategy): |
| 1. Focusing more on the moving hand |
| 2. Focusing less on the non-moving hand |
| 3. Physically relaxing the non-moving hand |
| 4. Increasing the rate of movement |
| 5. Increasing the force of the movement |
| 6. Increasing the size of the movement |
| 7. Tapping the fingers in a fixed sequence |
| 8. Tapping the fingers in a random sequence |
| 9. Opening and closing the hand |
| 10. Making grasping movements |
| 11. Imagining bilateral movements (while keeping the right hand still) |
| 12. - Other strategies (score) |

**Supplementary Table 1. Debriefing Questionnaire.** Participants filled in a questionnaire after the neurofeedback session.

| Questionnaire | | | | | | | | |
|---|---|---|---|---|---|---|---|---|
| A. How much control over the blue bar did you feel you had? (1-not in control, 5-full in control) | NF Group | | | | Sham Group | | | |
| | Association | | Dissociation | | Association | | Dissociation | |
| | Mean | SD | Mean | SD | Mean | SD | Mean | SD |
| | 2.63 | 1.2 | 3.1 | 0.98 | 2.70 | 0.59 | 2.95 | 0.64 |

**Supplementary Table 2. Debriefing Questionnaire Results.** There were no significant differences between sham and NF groups in response to the question "how much control over the blue bar did you feel you had?" for the Association condition (Mann-Whitney U test; $Z = 0.327$, $p=0.756$) or the Dissociation condition (Mann-Whitney U test; $Z = -1.081$, $p=0.314$). Within group, there were no significant differences in response to this question between experimental conditions (Wilcoxon signed-rank test; real NF group: $Z = -1.294$, $p=0.196$; Sham group: $Z = -0.991$, $p=0.322$). With regards to participants' rankings on how useful they found the strategies they tried (Supplementary Table 1, Question B), there was no difference in the rankings between groups for the Association condition (Mann-Whitney U test; $Z = 0.986$, $p = 0.349$) or the Dissociation condition (Mann-Whitney U test; $Z = 0.458$, $p = 0.654$). Overall, participants of both groups perceived similar degrees of control and considered that the strategies were similarly successful.

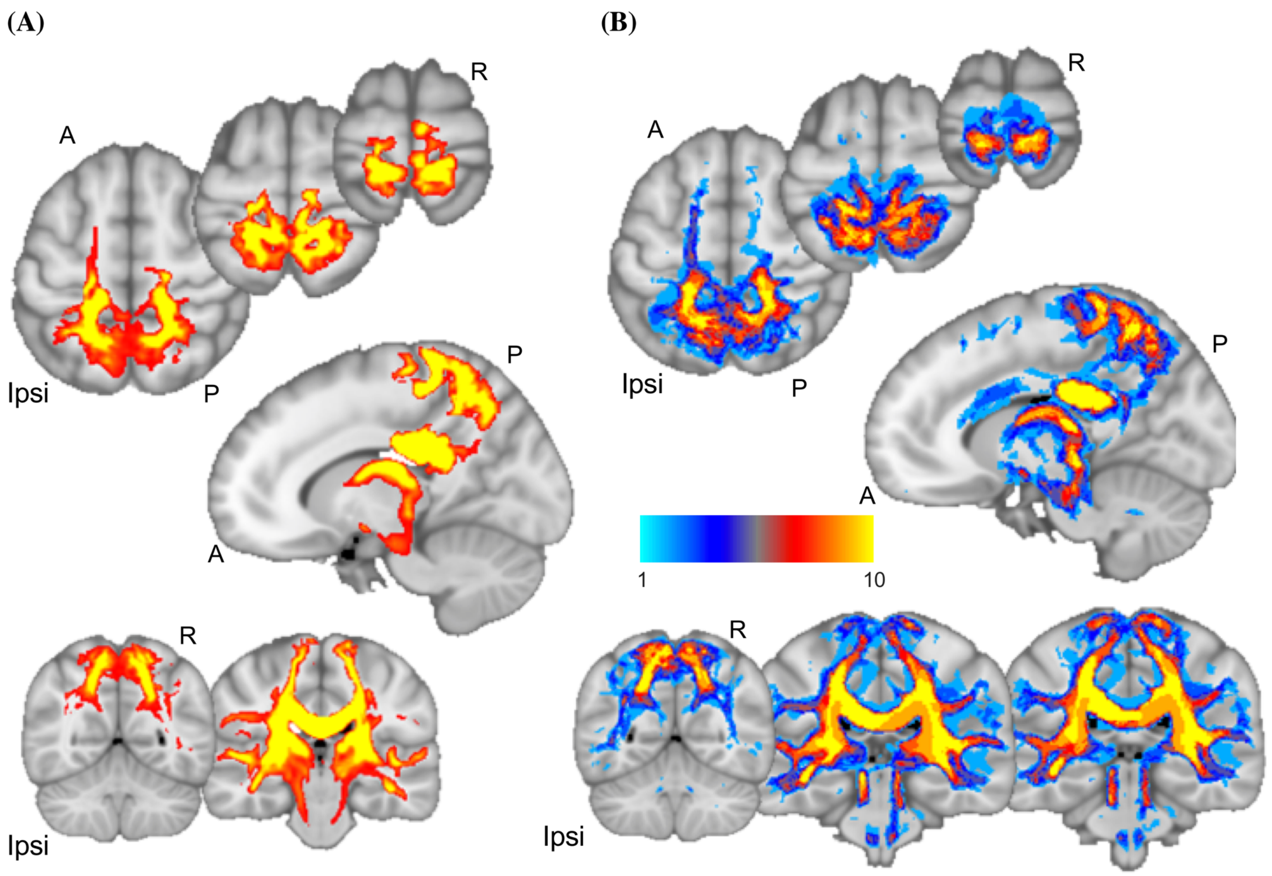


**Supplementary Figure 2. Significant FA cluster connects to sensorimotor and posterior parietal areas.** (A) Mean connectivity map (red-yellow) of all participants of the FA cluster. Yellow areas depict higher connectivity probability (threshold > 100). (B) Population connectivity map showing the overlap between participants (Light blue represents 1 participant – Yellow represents tracts common to the 10 participants). Tractography map is overlaid on the MNI template. Ipsi – Ipsilateral Hemisphere, A-Anterior, P- Posterior, R - Right.

| Participant | Session | Condition |
|---|---|---|
| 1 | 1 | Association |
| 2 | 2 | Dissociation |
| 3 | 2 | Dissociation |
| 4 | 1 | Dissociation |
| 5 | 1 | Association |
| 6 | 2 | Association |
| 7 | 1 | Dissociation |
| 8 | 2 | Dissociation |
| 9 | 1 | Dissociation |
| 10 | 2 | Association |

**Supplementary Table 3.** Session and condition of best performance for each participant.
